# Diurnal.plant.tools in 2024: expanding to *Marchantia polymorpha* and four angiosperms

**DOI:** 10.1101/2024.04.30.591826

**Authors:** Qiao Wen Tan, Emmanuel Tan, Marek Mutwil

## Abstract

Diurnal gene expression is a pervasive phenomenon occurring across all kingdoms of life, orchestrating adaptive responses to daily environmental fluctuations and thus enhancing organismal fitness. Our understanding of the plant circadian clock is primarily derived from studies in Arabidopsis and direct comparisons are difficult due to differences in gene family sizes. To this end, the identification of functional orthologs based on diurnal and tissue expression is necessary. The diurnal.plant.tools database constitutes a repository of gene expression profiles from 17 members of the Archaeplastida lineage, with built-in tools facilitating cross-species comparisons. In this database update, we expand the dataset with diurnal gene expression from 4 agriculturally significant crop species and Marchantia, a plant of evolutionary significance. Notably, the inclusion of diurnal gene expression data for Marchantia enables researchers to glean insights into the evolutionary trajectories of the circadian clock and other biological processes spanning from algae to angiosperms. Moreover, by integrating data from tissue, and perturbation gene expression datasets for Marchantia from other databases, the diurnal dataset complements it to offer a comprehensive overview of gene functions across varied biological contexts. This expanded database serves as a valuable resource for elucidating the intricacies of diurnal gene regulation and its evolutionary underpinnings in plant biology.

## Introduction

Diurnal gene expression is pervasive across all domains of life, from single-cellular bacteria to multicellular organisms such as plants and animals. The rhythmic pattern of gene expression in an approximately 24-hour cycle allows organisms to anticipate and respond efficiently to daily changes in their environment, thus allowing them to optimise their use of resources and fitness (Greenham and McClung, 2015). Diurnal expression of genes is driven by endogenous and non-endogenous oscillations, which are governed by the circadian clock and the integration of environmental responses respectively (Sanchez and Kay, 2016). In photosynthesising organisms, light is one of the most crucial environmental cues that entrains the circadian clock (Hsu and Harmer, 2014). Together, the endogenous and non-endogenous drivers regulate gene expression through molecular mechanisms such as splicing, polyadenylation (Yang *et al*., 2020) and protein-protein interaction (Hsu and Harmer, 2014).

Given the importance of photosynthesis which plants rely on to obtain energy, it is not surprising that the quality and amount of light sensed by the photoreceptors affects photosynthesis and related processes such as carbon metabolism, growth and response to stresses (Hsu and Harmer, 2014). Developmental processes such as flowering are also dependent on day length to ensure that such costly developmental processes occur at the right time of the year to avoid harsh weather and pollination success for maximal reproductive fitness (Laosuntisuk, Elorriaga and Doherty, 2023). Anticipatory defence mechanisms, such as the accumulation of jasmonic acid during the day when herbivores are active allow the plant to channel limited resources to the most important processes in a timely manner (Greenham and McClung, 2015).

The circadian clock is also an important consideration when studying stress responses. Importantly, genes involved in the cold stress response, particularly *COLD-REGULATED GENE 27* (*COR27*)/*COR28* are found to be part of the circadian clock component (Wang *et al*., 2017). In turn, cold-responsive genes are enriched for cis-regulatory motifs such as the *CCA1* binding site and the evening element which are recognised by *CCA1*, *LHY* and *REVEILLE* clock genes (Mikkelsen and Thomashow, 2009). Thus, a thorough understanding of how circadian genes are involved in various biological processes and stresses can affect how we tackle pressing issues such as the selection and breeding of plants under a rapidly changing climate (Laosuntisuk, Elorriaga and Doherty, 2023).

In a comprehensive review of the circadian clock across the plant kingdom, it was observed that the components were largely conserved across the plant kingdom, but with increasing copies of the genes in higher plants, and not all orthologs are rhythmically expressed (Laosuntisuk, Elorriaga and Doherty, 2023). This indicates the complexity of doing cross-species comparison and inference of the circadian clock especially over large evolutionary distances. Moreover, the increased number of clock genes indicates the genetic capacity of higher plants to have more fine-tuned control of the circadian rhythm. In order to gain a better understanding of the functions of circadian clock genes, consideration of the spatial temporal expression of the genes should also be considered.

The amount of diurnal gene expression data is limited especially for understudied species. Since the launch of the diurnal.plant.tools database which contained diurnal gene expression data for 17 species across Archaeplastida, diurnal gene expression data for four additional species, *Brassica rapa* (Kim *et al*., 2019)*, Hordeum vulgare, Setaria italica* (Lai *et al*., 2020) and *Sorghum bicolor* (Müller *et al*., 2020), which are important crop plants, are now available. While the expression of circadian clock genes has been studied in *Marchantia polymorpha* (Linde *et al*., 2017), there has been no update in a comprehensive diurnal gene expression dataset for this basal land plant that has now become an important model for evolutionary studies.

Gene co-expression analyses has been shown to be useful in the elucidation of genes that are involved in similar biological functions (Van Dam *et al*., 2017). Moreover, cross-species comparison of gene co-expression modules can increase the confidence of often noisy co-expression data when components of a gene module are observed across species (Proost, Krawczyk and Mutwil, 2017; Julca, Tan and Mutwil, 2023). Through the provision of gene co-expression data and a suite of tools for the elucidation of co-expressed genes, gene modules and comparative analyses, we envision the database to be an important hypothesis generation tool for biologists with 22 species in Archaeplastida.

## Methods

### Growth and experimental conditions for *Marchantia polymorpha*

Gemmae of male *Marchantia polymorpha* Takaragaike-1 were plated on half-strength Gamborg B-G Basal agar (0% sucrose, pH 5.5) in triplicates for each sampling time point. The cultures were maintained at 24°C under continuous LED light at 60 μEm^-2^s^-1^ for 8 days. On day 8, the cultures were transferred to a plant growth chamber (HiPoint M-313) and grown at 24°C under LED light at 60 μEm^-2^s^-1^ with a 12 h/12 h day/night cycle. Plants (3 plants in 2 mL Eppendorf tubes) were harvested at Zeitgeber time (ZT) 2, 6, 10, 14, 18, and 22 on day 15 and quenched immediately in liquid nitrogen and stored at -80°C until ready for RNA extraction.

### RNA isolation and sequencing

Each replicate was ground into a fine powder using a mortar and pestle with liquid nitrogen. Approximately 0.1g of grounded plant material was used for total RNA extraction using Protocol A (750 μL Binding Solution) of the SpectrumTM Plant Total RNA Kit (Sigma, STRN-250) according to the manufacturer’s instructions. Additionally, on-column DNase digestion was performed using 60 μL of DNase mixture (15 μL RQ1 RNase-Free DNase (Promega, M6101), 6 μL RQ1 DNase 10X Reaction Buffer and 19 μL nuclease-free water) per column.

An estimation of the quality and quantity of RNA was determined using Nanodrop before the samples were handed to Novogene (Singapore) for further quality control, library preparation and sequencing. Sample quantitation, integrity and purity were determined using Nanodrop, agarose gel electrophoresis and Agilent 2100 Bioanalyzer. Novogene (Singapore) conducted library construction from total RNA using the NEBNext® Ultra™ II Directional RNA Library Prep Kit for Illumina®, which involved enriching eukaryotic mRNA using oligo(dT) beads, selecting library sizes, and amplification via PCR. Sequencing was performed in paired-end mode with 150 base pairs, targeting a sequencing depth of approximately 20 million reads per sample on the Illumina Novaseq-6000 platform.

### Downloading publicly available diurnal transcriptomic data

Publicly available diurnal gene expression data for four species that were not included in the diurnal.plant.tools database (Ng *et al*., 2020) were downloaded from ENA (Leinonen *et al*., 2011)using the LSTrAP-Cloud pipeline (Tan, Goh and Mutwil, 2020). A total of 12, 18, 24 and 24 experiments were downloaded for *Brassica rapa* (Kim *et al*., 2019), *Sorghum bicolor*, *Setaria italica* (Lai *et al*., 2020) and *Hordeum vulgare* (Müller *et al*., 2020) respectively. Details on the experimental setup, sampling time points, number of replicates and BioProject IDs are reflected in Supplementary Table 1. The RNA-seq experiments were annotated using the metadata downloaded earlier from SRA with additional information from the publications.

### Quantification of gene expression

Coding sequences (CDS) of each species were downloaded from the corresponding sources in Supplementary Table 2 and used to create an index using the ‘*index’* command of the kallisto v0.46.0 software (Bray *et al*., 2016). Gene expression of the RNA-seq data was quantified using the ‘*quant*’ command of the kallisto v0.46.0 (Bray *et al*., 2016) software with the following parameters “–single -l 200 -s 20 -t 2” (single end experiment, average fragment length of 200 base pairs (bp) with a standard deviation of 20, and to run with two threads). The transcripts per million (TPM) values were then used to generate a gene expression matrix for each species.

### Quality control of publicly available RNA-seq data

The number and percentage of pseudoaligned reads were obtained for each RNA-seq experiment from the Kallisto output files to evaluate the quality of the RNA-seq experiment. Scatterplots of the percentage of pseudoaligned reads against the number of pseudoaligned reads were plotted for each species. Thresholds of at least 30% or 1,000,000 pseudoaligned reads were chosen, and the experiments that passed were used for downstream analyses.

### Detection of rhythmic genes

Rhythmic genes for *Brassica rapa, Hordeum vulgare, Setaria italica*, *Sorghum bicolor* and *Marchantia polymorpha* were identified using JTK_Cycle version 3.1 (Hughes, Hogenesch and Kornacker, 2010, Supplementary Tables S3-S7). Timepoints ZT4 and ZT20 for *Brassica rapa* and ZT2 and ZT14 were omitted for *Hordeum Vulgare* to ensure even time points. *Sorghum bicolor* and *Setaria italica* lacked replicates but were sampled for 72 hours. Thus, samples that corresponded to the same time in the 24-hour cycle were treated as replicates. The parameters used for each of the species can be found in Supplementary Table 8. Genes are considered to be expressed if the TPM value is greater than 1 in at least one replicate; additionally, genes are considered rhythmic if supported by a p-value < 0.05.

The JTK_Cycle algorithm treats the first sample as ZT0, but some species’ first sampling occurred at different times: *Sorghum bicolor* and *Setaria italica* at ZT3 or three hours after initial light exposure, and *Marchantia polymorpha* at ZT2 or two hours after initial light exposure. Thus, the LAG value predicted by JTK_Cycle was corrected by three hours for *Sorghum bicolor* and *Setaria italica* and for two hours for *Marchantia polymorpha*. For each species, a heatmap of normalised Z-score values of the significantly rhythmic genes TPM values was generated to visualise the overall diurnal expression profile (Supplementary Figure 1).

### Comparison of diurnal gene expression in *Marchantia* to other species

We compared the diurnal expression of 1-to-1 Marchantia orthologs to other species of the plant kingdom as done in a previous study (Ferrari *et al*., 2019). JTK outputs of *Cyanophora paradoxa, Chlamydomonas reinhardtii, Porphyridium purpureum, Klebsormidium nitens, Physcomitrella patens, Selaginella moellendorffii, Picea abies, Oryza sativa,* and *Arabidopsis thaliana* were retrieved from the kingdom-wide diurnal gene expression study (Ferrari *et al*., 2019, Supplementary Table 9-17). In addition, we also looked at the number of orthologs that have the same phase in *Marchantia* and another species after applying a phase shift of the observed LAG in the other species.

### Processing of data for update of database

Data for five new species (*Brassica rapa, Hordeum vulgare, Marchantia polymorpha, Setaria italica* and *Sorghum bicolor*) were formatted to update the diurnal.plant.tools database (http://diurnal.plant.tools/), an implementation of the CoNekT framework (Proost and Mutwil, 2018). CDS sequences of the respective species were uploaded to the database as described in Supplementary Table 2. The protein sequences available on the database are translated based on the CDS. Gene annotation for each species was completed by uploading each species’ fasta sequences to Mercator 4 version 5, an online tool for the functional annotation of land plant species (Schwacke *et al*., 2019). For each species, protein domain and Gene Ontology (GO) term were predicted using Interproscan version 5.59-91.0 (Jones *et al*., 2014). The parameters “-appl Pfam -goterms” were applied to utilise the Pfam protein database and include the GO Terms in the output file. Orthogroups were identified and obtained by inputting the protein sequences of all the existing species in the database and those of the new species into Orthofinder v.2.3.1 (Emms and Kelly, 2015). Gene annotations, GO terms, Interproscan results, and Orthofinder outputs (orthogroups and gene trees) were uploaded to supplement the CoNekT database. Annotation of the experiments, time of the day and diurnal time points were uploaded as “Condition specificity”, “Tissue specificity” and “ZT time” respectively (Supplementary Table 18).

The Pearson Correlation Coefficient (PCC) of all gene pairs in the gene expression matrix was calculated for each species using the script pcc.py from the Large Scale Transcriptome Analysis Pipeline (LSTrAP) Github (Proost, Krawczyk and Mutwil, 2017) and uploaded to the database. The database framework constructs the co-expression network by calculating the highest reciprocal rank (HRR) between gene pairs with a threshold of <= 100 (Proost and Mutwil, 2018). Subsequently, co-expression gene clusters were identified employing the heuristic cluster chiseling algorithm (Mutwil *et al*., 2009).

### Rhythmicity and co-expression of biological processes

To visualise the co-expression and rhythmicity of biological processes in *Marchantia polymorpha*, significantly rhythmic genes were filtered for the selected Mapman bins (Supplementary Table 19), which were obtained from the Mercator 4 v2.0 (Schwacke *et al*., 2019) with default parameters and MpTak1 v5.1R1 protein sequences (Montgomery *et al*., 2020) as input. The significantly rhythmic genes of the biological processes mentioned were used as input in the “Create custom network” tool with default parameters and *Marchantia polymorpha* selected as the species and network. to generate a custom co-expression network using the updated diurnal.plant.tools database. The JSON file of the network was downloaded from the results page and processed in Cytoscape version 3.10.1 (Shannon *et al*., 2003, Supplementary Data 1).

## Results

### Establishment of five new species in diurnal.plant.tools

Here, we provide an update of our diurnal gene expression database with data from 5 new species - *Brassica rapa, Hordeum vulgare, Setaria italica* and *Sorghum bicolor*, which were obtained from published studies and *Marchantia polymorpha*. Due to the lack of diurnal gene expression data for a basal land plant, we entrained the plant to a 12-hour light / 12-hour dark cycle and sampled at 6 time points every four hours, starting from ZT2 (2 hours after light). The quality of experiments and genes was evaluated through PCA analyses (Figure 1A and B), and we observe that the variation of the experiments and genes reflects the order of the diurnal timepoint as expected. Experiments corresponding to the same biological replicate or timepoint tend to cluster closer to each other than to other replicates and timepoints.

The distribution of diurnally rhythmic genes was visualised across different phases in a 24-hour period (Figure 1C). Distribution of the diurnally rhythmic genes was either unimodal, as seen in *H. vulgare* (ZT23), *S. bicolor* (ZT16) and *S. italica* (ZT10-ZT16) respectively or bimodal, as seen in *B. rapa* (ZT0 and ZT12) and *M. polymorpha* (ZT10 and ZT20).

**Figure 1.**
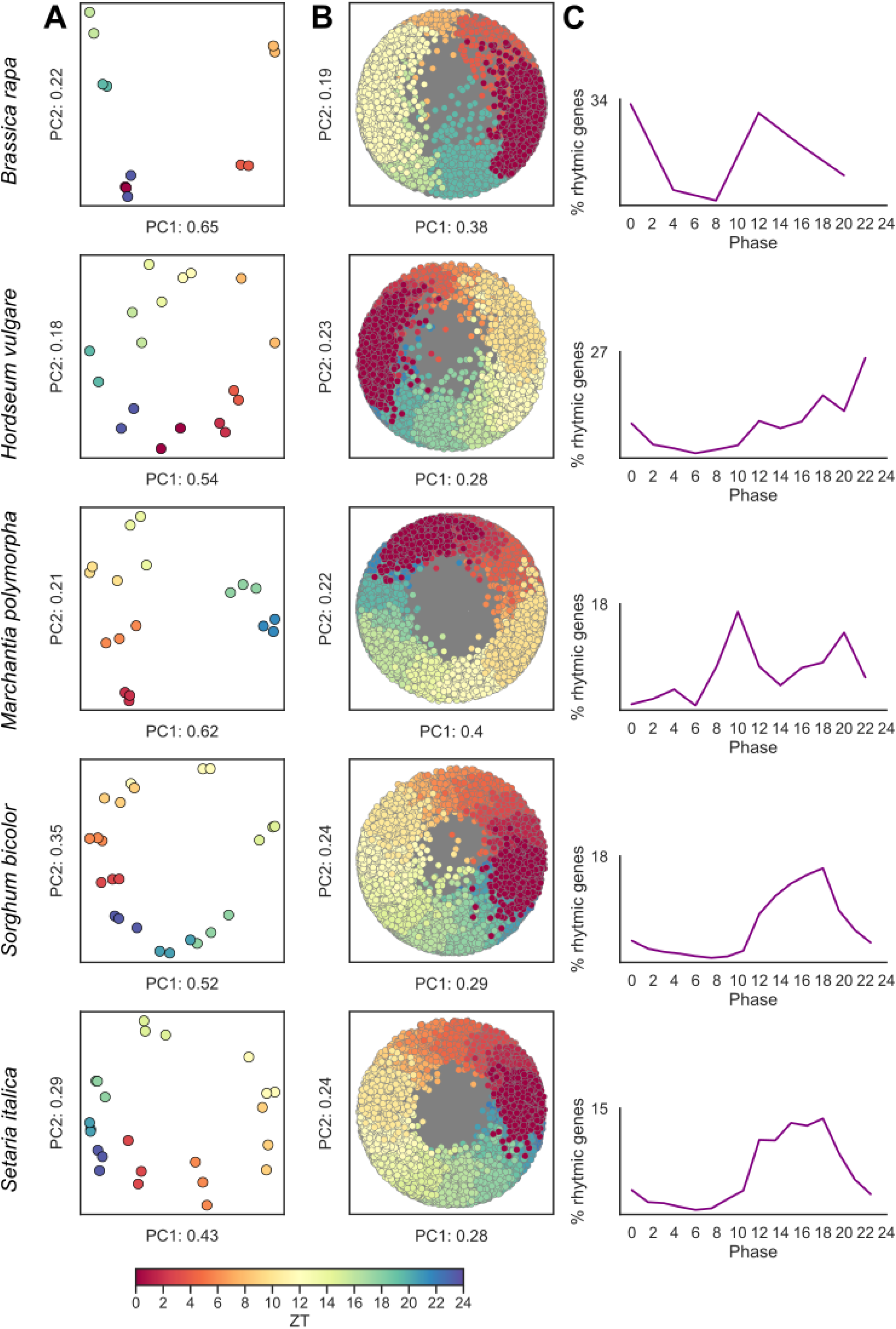
Overview of diurnal gene expression data. A) Principal Component Analysis (PCA) of RNA-sequencing experiments coloured according to Zeitgeber (ZT) time points. B) PCA of genes expression profile coloured according to the phase identified by JTK_Cycle (grey points are non-rhythmic genes). C) Distribution of rhythmic genes (as %, y-axis) across the phases identified by JTK_Cycle (x-axis).

### Comparative analysis reveals the conservation of diurnal gene expression between Marchantia and other Archaeplastida

The normalised gene expression of individual RNA-sequencing experiments across 6 timepoints (ZT2, 6, 10, 14, 18 and 22) in triplicates showed an expected phasing pattern (Figure 2A). We identified 51.1% of genes with significantly rhythmic gene expression (adjusted p-value < 0.05) with Jonckheere-Terpstra-Kendall (JTK_Cycle) algorithm, which revealed the time of the day where each rhythmic gene peaked (phase, Supplementary Table 7). The distribution of phases showed two peaks at ZT10 (i.e., 10 hours after lights on) and ZT20 (i.e., 4 hours before lights on, Figure 6B), which was reminiscent of phase distribution in Arabidopsis and rice (Ferrari *et al*., 2019). Our previous study indicated that cell division genes are peaking just before the light / dark change in single cellular algae *Cyanophora paradoxa*, *Porphyridium purpureum*, *Chlamydomonas reinhardtii* and simple multicellular algae *Klebsormidium nitens* (Ferrari *et al*., 2019), showing that the diurnal rhythm regulates cell division in these algae. To elucidate whether cell cycle genes are also under strong diurnal control in Marchantia, we looked at the normalised gene expression of cyclin A, B, D, cyclin-dependent kinases (CDKs), DNA primase and DNA polymerase across the 6 sampling time points (Figure 2C). In contrast to the single cellular algae, Marchantia showed a more uniform expression of cell cycle genes (Figure 2C), indicating that, similarly to other land plants, Marchantia’s cell cycle is not under the control of diurnal rhythm.

To elucidate how the different biological pathways are active during a typical day of Marchantia, we visualized the phase distribution of rhythmic genes and their pathways and performed a clustering analysis to reveal which pathways show similar diurnal gene expression (Figure 2D). Not surprisingly, we observed that, e.g., photosynthesis genes peak in expression during the day (Figure 2D, photosynthesis is labelled green during ZT 0-10) and shows a similar pattern with redox homeostasis, which is important for scavenging harmful reactive oxygen species generated by light capture (Das *et al*., 2015). Interestingly, and in line with the phase distribution plot (Figure 2A), we observed that most biological pathways tend to peak twice a day: during the day at ∼ZT10 and the night at ∼ZT20.

Our previous study showed that diurnal gene expression is conserved across >1 billion years of plant evolution (Ferrari *et al*., 2019). To investigate whether Marchantia’s diurnal gene expression is also conserved, we compared the distribution of phases of Marchantia genes and their orthologs in four algae (*Cyanophora paradoxa*, *Porphyridium purpureum*, *Chlamydomonas reinhardtii,* and *Klebsormidium nitens*) and five land plants (*Physcomitrella patens, Selaginella moellendorffii, Picea abies, Oryza sativa,* and *Arabidopsis thaliana*).

**Figure 2.**
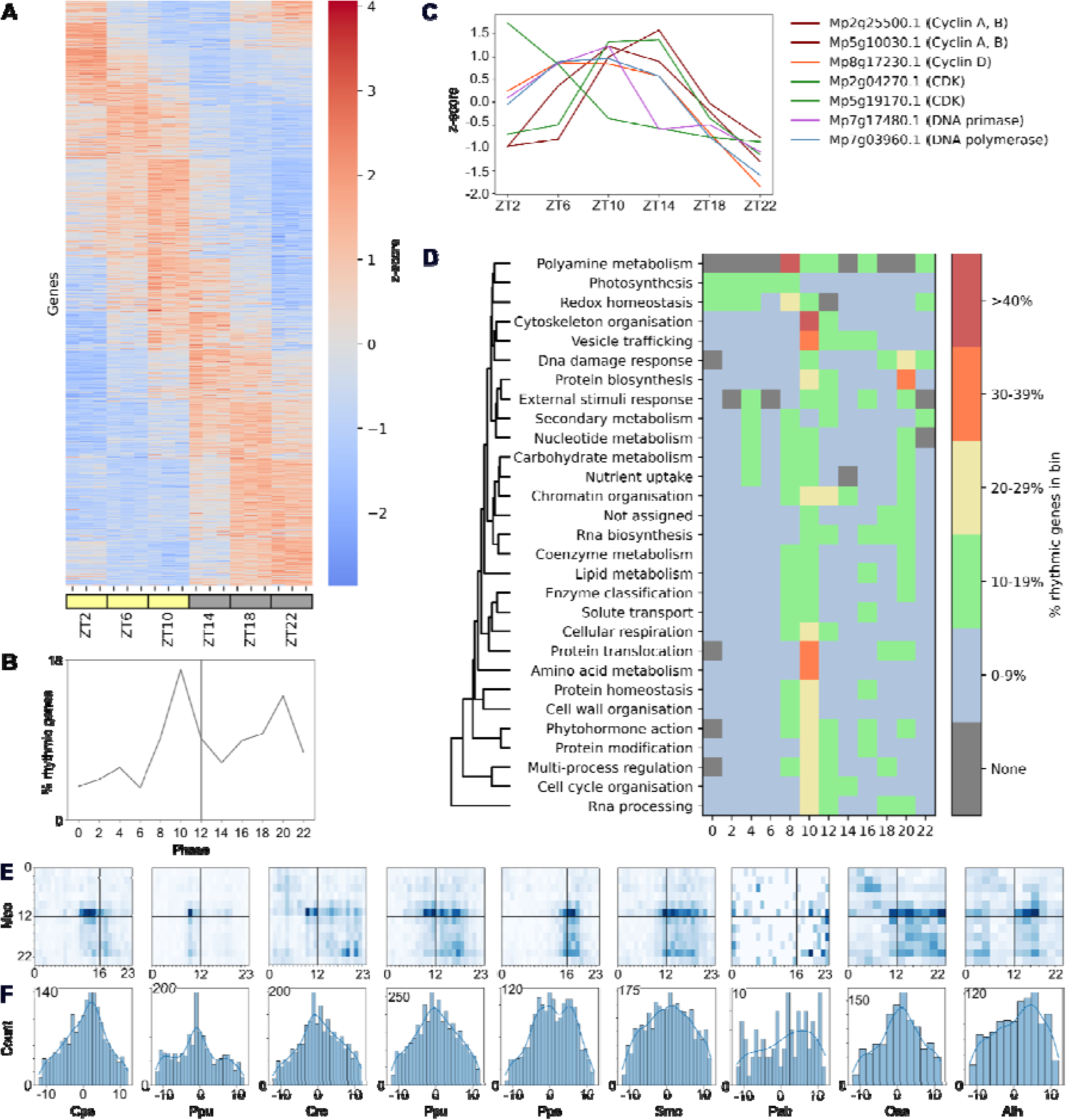
Comparison of diurnal rhythms of Marchantia and other Archaeplastida. A) Z-score-normalised expression of genes (y-axis) taken at different Zeitgeber timepoints (x-axis). Yellow and grey boxes indicate the presence and absence of light at the time of sampling, respectively. The ticks indicate the biological replicates. The data was collected in triplicates. B) Distribution of rhythmic genes (as %, y-axis) across phases (x-axis). C) Z-score-normalised expression of significantly rhythmic (adjusted p-value < 0.05) cell cycle genes (cyclin A, B, D, cyclin-dependent kinase (CDK), DNA primase, and DNA polymerase). The y-axis and x-axis indicate the gene expression and average expression at a given ZT, respectively. D) Diurnal phase peak distribution of biological pathways. Rows represent biological pathways, while the columns indicate the phases at which the genes in a pathway peak. Cells are coloured according to the percentage of rhythmic genes in a given pathway at each phase which add up to 100%. The linkage between the Mapman bins was determined by scipy hierarchical clustering. E) The heatmap shows the phase distribution of phases of Marchantia genes (y-axis, phases are arranged from 0:top to 23:bottom) versus the phases of the 1-to-1 orthologs from other species (x-axis, phases are arranged from 0:left to 23:right). The intensity of the cells in the heatmap is proportional to the number of genes that peak at a given phase for the two species. Black lines indicate the light-to-dark transition. Abbreviations used to represent species are as follows: *Cyanophora paradoxa* (Cpa), *Chlamydomonas reinhardtii* (Cre), *Porphyridium purpureum* (Ppu), *Klebsormidium nitens* (Kni), *Physcomitrella patens* (Ppa), *Selaginella moellendorffii* (Smo), *Picea abies* (Pab), *Oryza sativa* (Osa) and *Arabidopsis thaliana* (Ath). F) Distribution of differences of phases of the 1-to-1 orthologues between Marchantia and the nine members of Archaeplastida. A value of 0 indicates that the 1-to-1 orthologs peak at the same time of the day; the values of -10 and 10 show that the orthologs peak 10 hours earlier and 10 hours later in other species than in Marchantia, respectively.

The analysis plots the phases of Marchantia genes (y-axis, phases are arranged from 0 (top) to 23 (bottom)) versus the phases of the 1-to-1 orthologues from other species (x-axis, phases are arranged from 0 (left) to 23 (right)). Two species with similar diurnal gene expression would show an accumulation of points at similar x and y coordinates, indicating that the orthologs of the two species peak at the same time of the day. Conversely, the points would be randomly distributed in the plot for two species with different diurnal gene expressions. The analysis revealed that for most species, with the exception of Selaginella and Picea, we observe groups of genes at similar x and y coordinates, suggesting that the diurnal gene expression of Marchantia is conserved (Figure 2E). To further quantify the conservation of diurnal gene expression, we plotted the distribution of the differences between phase values of one-to-one orthologues of Marchantia and the nine other species. For all species, except for Selaginella, Picea, and Arabidopsis, we observed a clear peak at around 0 (Figure 2F), confirming that the diurnal gene expression of Marchantia is conserved to most Archaeplastida. Taken together, the conservation between Marchantia and the other Archaeplastida makes Marchantia an attractive model to study diurnal gene expression.

### Inferring functional orthologs through diurnal and organ expression

The MYB domain-containing transcription factor gene family consist of genes that are key components of the circadian clock. All plants consist of at least one member in this gene family but differ in the representation in *CIRCADIAN CLOCK-ASSOCIATED 1* (*CCA1*) and *LCL/REVEILLE* (*RVE*) clades. In higher plants, there is a larger number of circadian clock components due to events such as genome duplication and polyploidization as compared to basal plants, and this is especially true for the MYB transcription factors (10 copies in Arabidopsis) and the *CCT* protein (17 copies in Arabidopsis) (Laosuntisuk, Elorriaga and Doherty, 2023). In order to effectively compare the circadian clocks across species, information regarding a gene’s spatial-temporal expression is necessary on top of sequence similarity.

The diurnal.plant.tools and conekt.plant.tools (Proost and Mutwil, 2018) databases have one of the most comprehensive collections of gene expression data for plants based on diurnal and tissue expression respectively. To demonstrate how the databases can give us more insights inferring functional orthologs, we first went to the profile of MpRVE (Mp2g14310, https://diurnal.sbs.ntu.edu.sg/sequence/view/1004422) and clicked on the link to the gene family tree (Supplementary Figure 2, https://diurnal.sbs.ntu.edu.sg/tree/view/154077)which displays the normalised expression profile of the genes across ZT groups (Supplementary Figure 2, https://diurnal.sbs.ntu.edu.sg/tree/view/154077). From the interactive tree, we could see that not all members are rhythmically expressed and 1 additional Marchantia gene, *Mp1g25720,* has not been annotated as an RVE gene and is not rhythmically expressed (Supplementary Table 7). To compare the spatial-temporal expression of Marchantia and Arabidopsis MYB transcription factor genes, we downloaded the table of 166 genes in the gene family (Supplementary Table 20). From the gene family, we selected the 10 Arabidopsis genes (*AT1G01520*, *AT1G18330*, *AT2G46830*, *AT3G09600*, *AT3G10113*, *AT4G01280*, *AT5G02840*, *AT5G17300*, *AT5G37260* and *AT5G52660*) and 2 Marchantia genes (*Mp2g14310* and *Mp1g25720*) as input to the create heatmap tool in diurnal.plant.tools (Figure 3A) and conekt.plant.tools respectively (Figure 3B). In alignment with what is known about the expression of circadian clock MYB transcription factors (Laosuntisuk, Elorriaga and Doherty, 2023), 8 of the 10 Arabidopsis orthologs are rhythmic (Supplementary Table 17).

The tissue expression of the MYB transcription factor on the other hand revealed organ-specific expression of the transcription factors. We hypothesise that the organ-specific transcription factors allow higher plants an opportunity to fine-tune the regulation of specific subsets of genes in various organs. While the expression of *AT1G01520*, *AT3G09600*, *AT4G01280* and *AT5G17300* peak at the same time as *MpRVE* (ZT4), they are unlikely to be functional orthologs as they are specifically expressed in the seeds, root, stem and root/flower respectively. Taking the diurnal and tissue expression of the MYB transcription factors, we can narrow down the functional orthologs of *MpRVE* to *AT2G46830* and *AT5G02840* which have peak expressions at ZT6 and ZT2 respectively. In addition, when sequence similarity is taken into consideration, *AT5G02840*, annotated as *RVE4* on TAIR (The Arabidopsis Information Resource (TAIR), no date b) is the most likely functional ortholog to *MpRVE* as they belong to the same clade (Supplementary Figure 2). *AT2G46830*, on the other hand, is annotated as *CCA1* on TAIR (The Arabidopsis Information Resource (TAIR), no date a), which has been noted to be lost in Marchantia (Laosuntisuk, Elorriaga and Doherty, 2023).

### Global analysis of the rhythmicity of biological pathways with diurnal.tools

Many biological processes follow a diurnal rhythm. Through the custom network tool on the database, we can easily gain an overview of the rhythmicity of the major biological process of interest by providing a list of genes.

**Figure 3.**
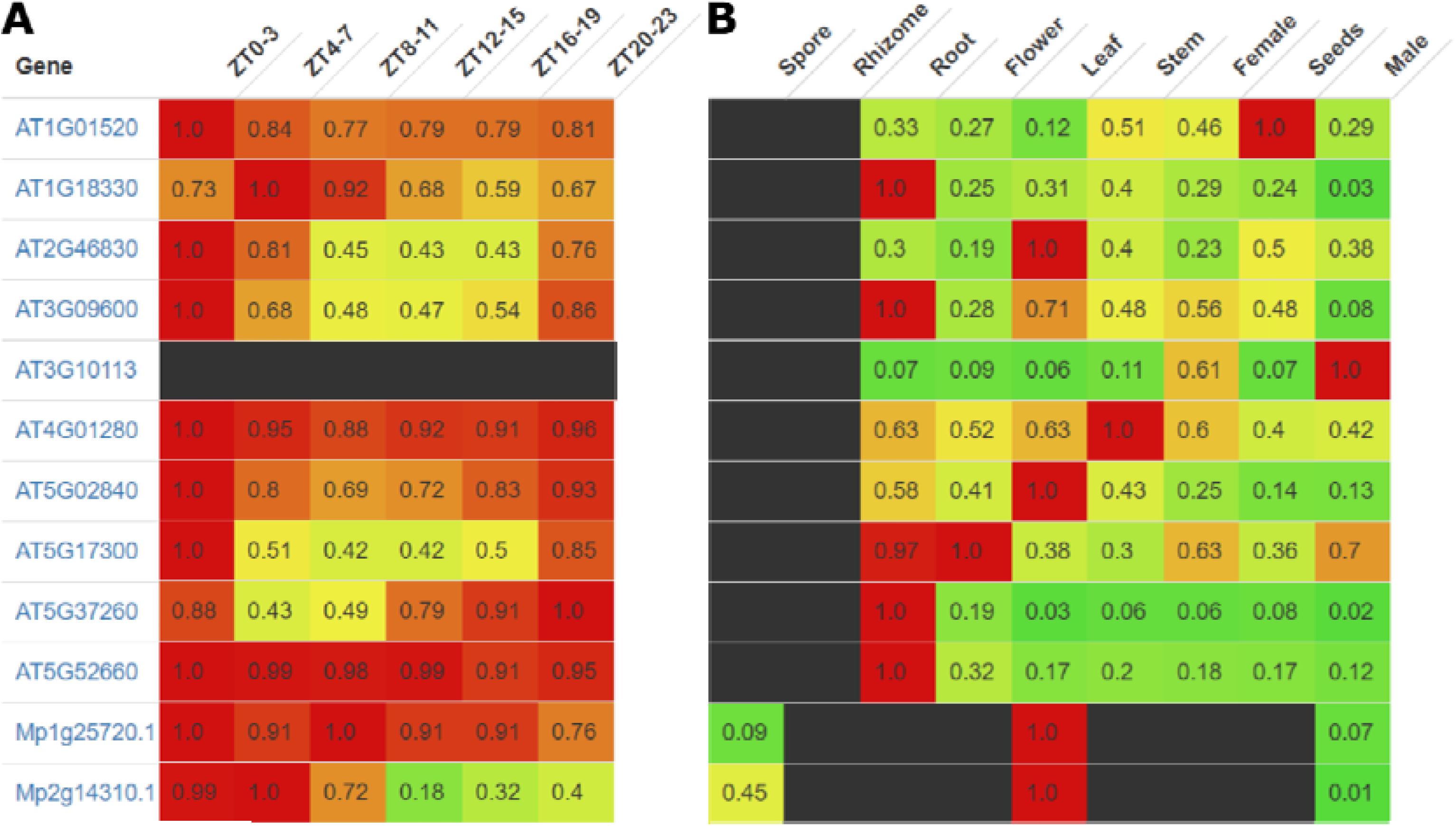
Expression of MYB domain transcription factor gene family in Marchantia and Arabidopsis. Row normalised expression of *RVE* gene family (OG_42_0000380 in diurnal.plant.tools) in A) diurnal.plant.tools B) conekt.plant.tools. Red indicates higher expression while green indicates lower expression.

The resulting co-expression network coloured by the time of peak expression of the genes reveals protein synthesis to be highly rhythmic but at different times of the day. Synthesis of proteins is concentrated in the plastid at the beginning of the day (ZT0-3) and switches to the mitochondria towards the middle of the day (ZT8-11) and finally to the cytoplasm at the end of the day (ZT16-23). Similar to plastidial protein synthesis, the expression of genes involved in photosynthesis typically peaks at ZT0-7, with some genes anticipating light, peaking at ZT20-23. Genes involved in starch and sucrose metabolism, processes that are tightly linked to photosynthesis also tend to peak at the start of the day (ZT0-3).

In the network, we noticed many edges between terpenoids and photosynthesis nodes. Upon further inspection of the genes connected to photosynthesis (Supplementary Data 1), the Mapman annotation of the genes revealed that they were involved in the methylerythritol phosphate (MEP) pathway and carotenoid biosynthesis. The strong association between genes of the MEP pathway, which generates precursors for carotenoid biosynthesis, is not surprising as carotenoids are essential light-harvesting pigments involved in photosynthesis and offer photoprotective and antioxidative effects against excess light and reactive oxygen species (Caferri *et al*., 2022). In addition, genes involved in the biosynthesis of flavonoids chalcones and aurones, and involved in the phenylpropanoid pathway are found to be correlated with carbohydrate metabolism frequently. The biosynthesis of flavonoids involves the shikimate pathway that converts erythrose 4-phosphate and phosphoenolpyruvic acid to phenylalanine which is then converted into 4-coumaroyl-CoA, a precursor of flavonoids, through a series of reaction in the phenylpropanoid pathway (Liga, Paul and Péter, 2023).

**Figure 4.**
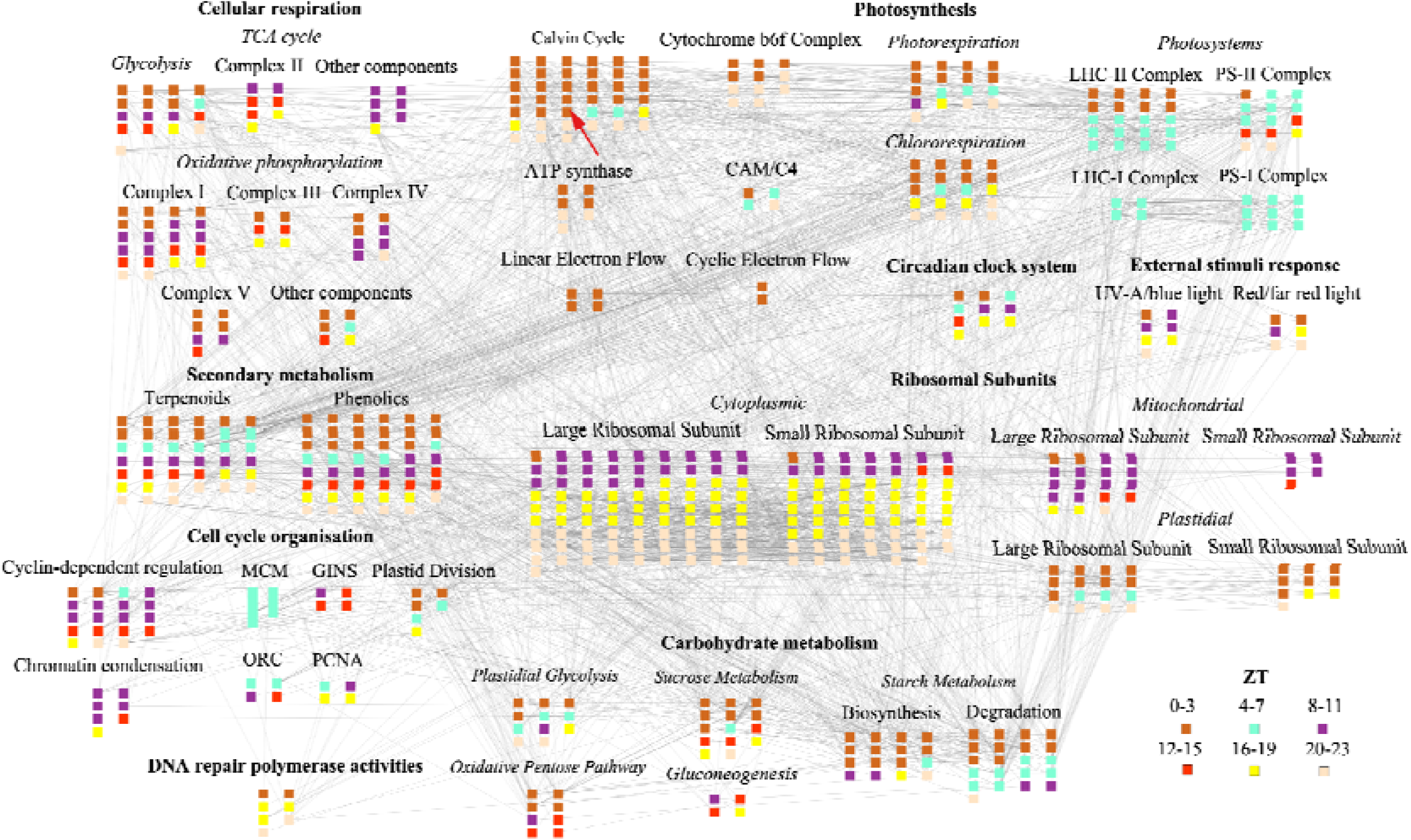
Custom co-expression network of core biological processes of *Marchantia polymorpha*. Nodes and edges represent genes and the co-expression relationship (HRR <= 100) between genes respectively. The time of peak gene expression is reflected in the node colours. Annotations in bold reflect the top-level Mapman bin while other annotations reflect finer bin annotations. Additional details regarding the bins selected are provided (Supplementary Table 19) and gene ID and other information can be found in the original Cytoscape file (Supplementary Data 1).

### Integration of Marchantia diurnal and stress data via diurnal.plant.tools and eFP browser

While gene expression data consisting of different plant organs or experimental perturbations are generally available for widely studied plant species, high-quality diurnal gene expression data is limited. Diurnal gene expression data provides a unique opportunity to examine the relationship between genes and biological processes under typical growth conditions within a specific organ or organism, depending on the sampling approach. In contrast, gene expression data from different tissues would underscore the similarities of these processes across various tissue types, while data from perturbations would illuminate how these processes respond to variable conditions.

Photosynthesis is an essential process for plants. From the network of core biological processes (Figure 4), we identified the most connected photosynthesis gene (i.e. gene with the most number of edges), *Mp4g06000*, a RUBisCO small subunit. The expression of *Mp4g06000* over different time points (Figure 5A), in different tissues and time points (Figure 5B) can be obtained from diurnal.plant.tools and *Marchantia* eFP (Tan *et al*., 2023) respectively. As expected, the expression of *Mp4g06000.1* peaks at the ZT2 sampling timepoint, in agreement with the corrected phase estimated by the JTK algorithm, reflected in Figure 4. Interestingly, the gene shows anticipatory behaviour where the expression increases as the night progresses (Figure 5A). While the eFP also contains the same diurnal gene expression in pictorial form, it is less informative in showing diurnal trends due to additional expression information from other datasets. However, the integration of gene expression levels in different organs and stress conditions in the eFP pictogram allows us to gain additional insights into where the gene is expressed and its response to a comprehensive set of single and stress combinations (Tan *et al*., 2023). We observe that *Mp4g06000.1* is most highly expressed in the thallus, the leaf equivalent organ of higher plants and the gene is lowly expressed (yellow cells) during dark stress while highly expressed (red cells) for light and light combination stresses, except for light-cold stress. In addition, osmotic stress induced by salt and mannitol moderately increases the expression of *Mp4g06000.1* (orange cells) and is especially high in the nitrogen-salt combination (red cells). The guilt-by-association principle suggests that co-expressed genes are likely to be involved in similar biological functions (Wolfe, Kohane and Butte, 2005). From the database, we retrieved the co-expression neighbourhood of *Mp4g06000.1* (https://diurnal.sbs.ntu.edu.sg/network/graph/565284) and downloaded the network in JSON format for further processing in Cytoscape v3.10.1 (Shannon *et al*., 2003) to gain a better understanding of what genes and processes it is connected to (Figure 4C). As expected, the 64 co-expression neighbours of *Mp4g06000* (Supplementary Table 21) are involved in photosynthesis (20 genes) or photosynthesis-related processes such as carbohydrate metabolism (6 genes) and secondary metabolism (2 genes). Surprisingly, the neighbourhood contained 27 genes that were not assigned to a Mapman bin. To investigate if we could infer the gene function from the gene family of the respective genes, we entered the gene information page on diurnal.plant.tools to access the gene family page and extracted the common annotation for the gene annotation, if any. Most unassigned genes (20) had some form of annotation that could be inferred from the gene family. Of which, orthologs of *Mp1g15330.1* (HVA22-like, Zhang *et al*., 2023), *Mp5g01770.1* (temperature-induced lipocalin, Abo-Ogiala *et al*., 2014) and *Mp5g15480.1* (interferon-related developmental regulator family protein, Park *et al*., 2009) have been implicated in stress response.

**Figure 5.**
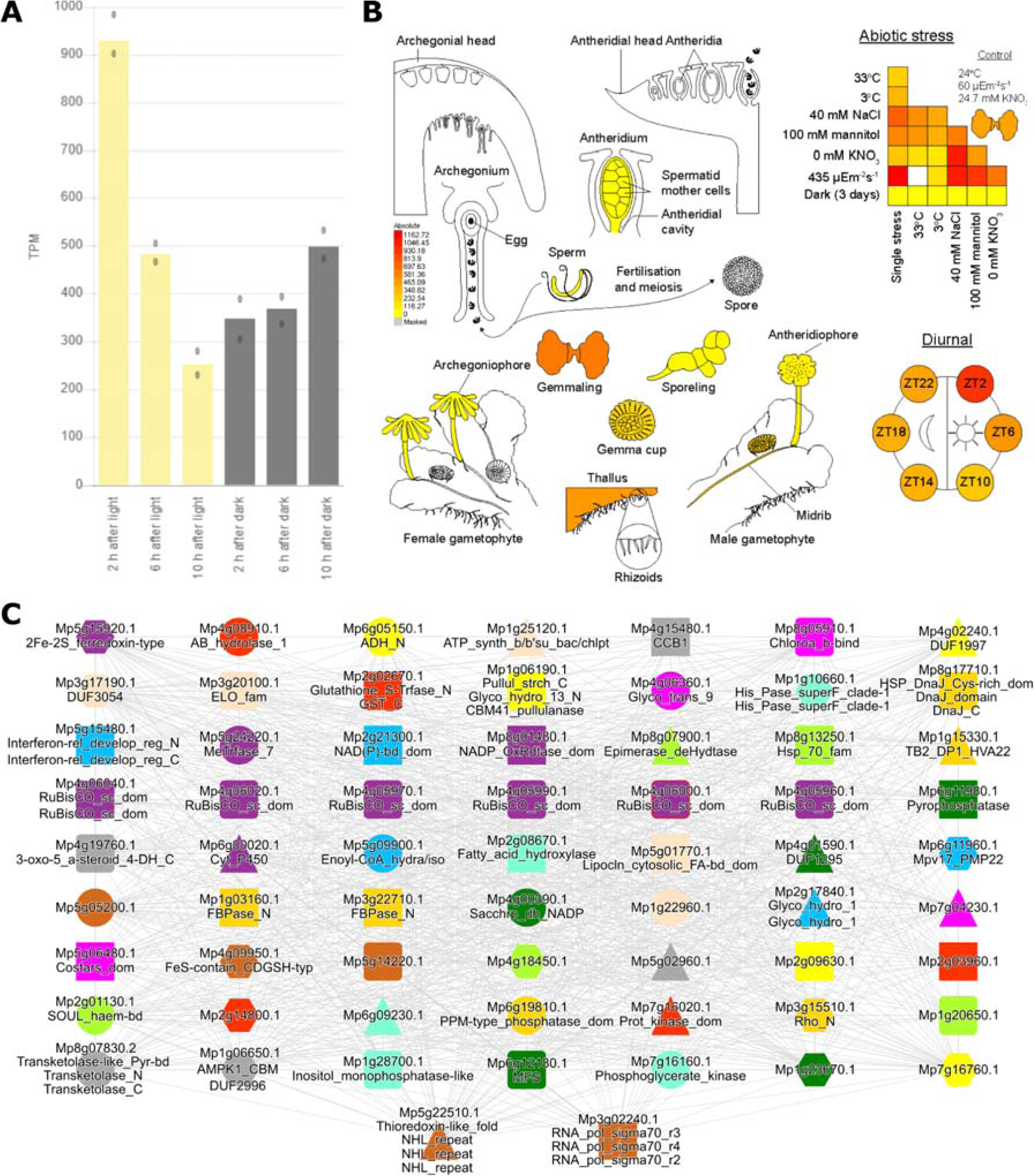
Expression of Mp4g06000.1 and co-expression neighbourhood. Expression profiles in A) diurnal.plant.tools B) eFP. C) Co-expression neighbourhood of *Mp4g06000.1* (red border). Unique node shapes and colours represent neighbours with common annotations (gene family/interpro). Gene identifiers are followed by interpro annotations. Nodes are connected by edges when the highest reciprocal rank is <= 100.

As some of the inferred gene annotations were associated with stress response, we hypothesise that this group of unassigned genes could contain genes that link the photosynthesis process to stress response. To this end, we observed 14 of these unassigned genes to be stress-responsive when viewed on eFP. This set of genes serves as prime candidates for further investigation on other factors that could influence photosynthesis and how photosynthesis could be regulated during stress response.

## Discussion

In plants, close to one-third of genes are diurnally expressed (Ferrari *et al*., 2019) through regulation by the circadian clock and entrainment by environmental cues such as light and temperature. The optimised expression of genes is not only necessary for the immediate response to environmental cues but also for the dynamic distribution of resources for the growth and development of the plant. Currently, our understanding of diurnal gene expression is based on what we know in Arabidopsis. However, organisms across the Archaeplastida lineage have differing amounts and types of known clock components (Laosuntisuk, Elorriaga and Doherty, 2023). In this database update, we expanded the number of species to include 4 more crop species and Marchantia, a basal land plant. Through the diurnal co-expression database, and other existing datasets for tissue expression and stress perturbations, such as conekt.plant.tools (Proost and Mutwil, 2018), eFP (Winter *et al*., 2007), researchers could refine their understanding of different circadian clock components across species by comparing the diurnal and tissue expression patterns to narrow down on functional orthologs. Additionally, together with the availability of co-expression datasets during stress, researchers can gain a comprehensive understanding of the functional neighbourhood of genes at different times of the day, tissues and stress perturbations.

## Supporting information

Supplemental Data (Cytoscape)

## Data availability

The RNA-seq data capturing the diurnal gene expression of *Marchantia polymorpha* are available from https://www.ebi.ac.uk/ena as EMTAB-11141 [https://www.ebi.ac.uk/biostudies/arrayexpress/studies/EMTAB-11141].

## Code availability

Python and bash scripts used to generate the intermediate files and figures in the paper are available at https://github.com/emmanuel-tan/diurnal.plant.tools-2024.

## Author contributions

M.M. conceived the project, supervised E.T. and Q.W.T., wrote the manuscript. Q.W.T. performed the diurnal experiment, generated the RNA material for RNA-sequencing, data analysis, wrote the manuscript and supervised E.T.. E.T. performed the downstream analysis, updated diurnal.plant.tools and wrote the manuscript.

---

**Supplementary Figure 1.**
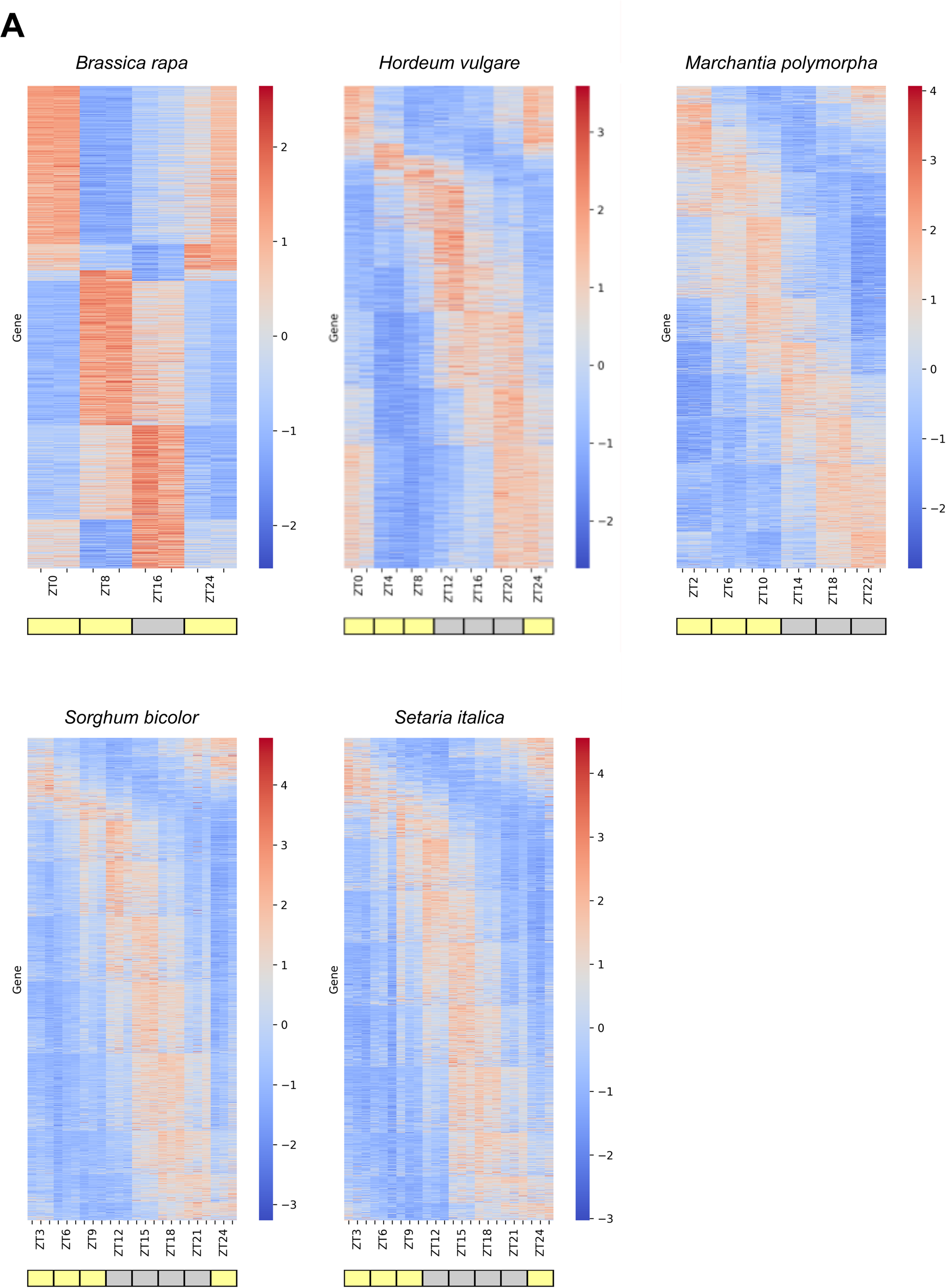
Heatmap of overall expression of diurnal genes. Each horizontal line represents a gene’s Z-score-normalised expression (y-axis) at various ZT timepoints (x-axis), while the colour indicates the expression level (higher and lower expression indicated by red and blue, respectively). Yellow and grey shading signify the presence and absence of light, respectively, during the sampling period, as per experimental conditions outlined. Biological replicates are marked along the axis with ticks.

Supplementary Figure 2. Phylogenetic tree of OG_42_0000380

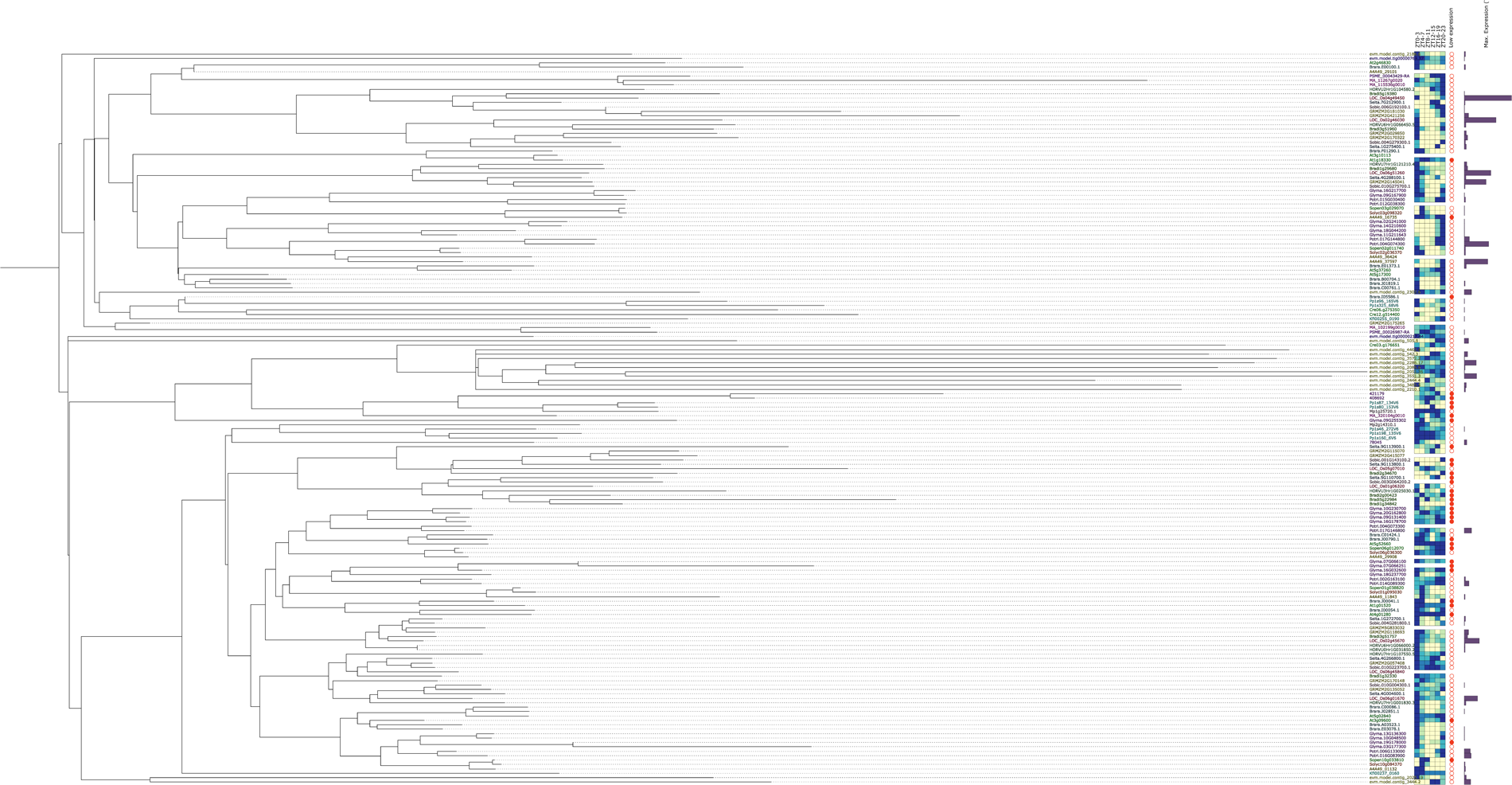

